# YeaZ: A convolutional neural network for highly accurate, label-free segmentation of yeast microscopy images

**DOI:** 10.1101/2020.05.11.082594

**Authors:** Nicola Dietler, Matthias Minder, Vojislav Gligorovski, Augoustina Maria Economou, Denis Alain Henri Lucien Joly, Ahmad Sadeghi, Chun Hei Michael Chan, Mateusz Koziński, Martin Weigert, Anne-Florence Bitbol, Sahand Jamal Rahi

## Abstract

The processing of microscopy images constitutes a bottleneck for large-scale experiments. A critical step is the establishment of cell borders (‘segmentation’), which is required for a range of applications such as growth or fluorescent reporter measurements. For the model organism budding yeast (*Saccharomyces cerevisiae*), a number of methods for segmentation exist. However, in experiments involving multiple cell cycles, stress, or various mutants, cells crowd or exhibit irregular visible features, which necessitate frequent manual corrections. Furthermore, budding events are visually subtle but important to detect. Convolutional neural networks (CNNs) have been successfully employed for a range of image processing applications. They require large, diverse training sets. Here, we present i) the first set of publicly available, high-quality segmented yeast images (>10’000 cells) including mutants, stressed cells, and time courses, ii) a corresponding U-Net-based CNN, iii) a Python-based graphical user interface (GUI) to efficiently use the system, and iv) a web application to test it (www.quantsysbio.com). A key feature is a cell-cell boundary test which avoids the need for additional input from fluorescent channels. A bipartite graph matching algorithm tracks cells in time with high reliability. Our network is highly accurate and outperforms existing methods on benchmark images recorded by others, suggesting it transfers well to other conditions. Furthermore, new buds are detected early with high reliability. We apply the system to detect differences in geometry between wild-type and cyclin mutant cells. Our results indicate that morphogenesis control occurs unexpectedly early in the cell cycle and is gradual, demonstrating how the efficient processing of large numbers of cells uncovers new biology. Our system can serve as a resource to the community, expanded continuously with new images. Furthermore, the techniques we develop here are likely to be useful for other organisms as well.

The identification of cell borders (‘segmentation’) in microscopy images constitutes a bottleneck for large-scale experiments. For the model organism *Saccharomyces cerevisiae*, current segmentation methods face challenges when cells bud, crowd, or exhibit irregular features. Here, we present i) the first set of publicly available, high-quality segmented yeast images (>10’000 cells) including mutants, stressed cells, and time courses, ii) a corresponding convolutional neural network (CNN), iii) a graphical user interface and a web application (www.quantsysbio.com) to efficiently employ, test, and expand the system. A key feature is a cell-cell boundary test which avoids the need for fluorescent markers. Our CNN is highly accurate, including for buds, and outperforms existing methods on benchmark images, indicating it transfers well to other conditions. To demonstrate how efficient, large-scale image processing uncovers new biology, we analyzed the geometries of ≈2200 wild-type and cyclin mutant cells and found that morphogenesis control occurs unexpectedly early and gradually.

## Introduction

Budding yeast is an important model organisms in genetics, molecular biology, systems biology, and synthetic biology. Almost all current segmentation methods for yeast images^1–8^ rely on classical image processing techniques^9^ such as thresholding, edge detection, contour fitting, and watershed. However, for many experiments, the segmentations produced by these tools require frequent user interventions. Common challenges for yeast image segmentation include (Fig. 1):

**Fig. 1:**
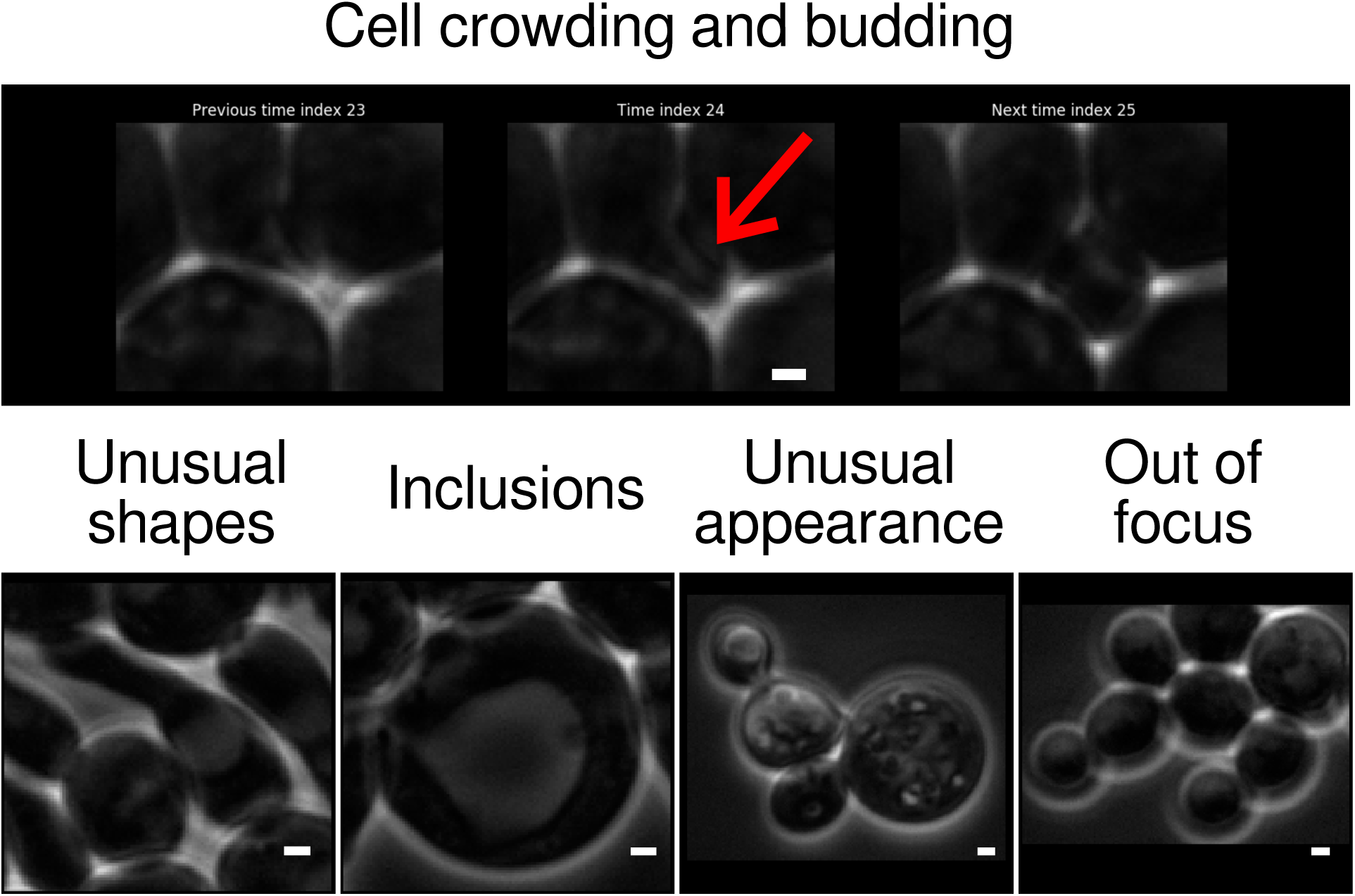
Challenging cases for the segmentation of yeast images. The red arrow points to a difficult-to-see new bud that appears in a timelapse movie. Scale bar: 1 *μ*m.

- cell crowding,
- irregular shapes,
- transparent inclusions (e.g., vacuoles),
- unusual visible features,
- budding events, and
- imperfect focus during imaging.

CNNs have established themselves in recent years as efficient and powerful computational models for segmentation tasks.^10^ CNNs replace sophisticated classical image processing algorithms with neural-network based models which are trained on a sufficiently large and diverse set of examples.^11^ A key advantage of CNNs over non-learning based approaches is that in order to improve the predictions for new cases or conditions, fundamentally new ideas are not needed. In principle, new cells or conditions that the system performs poorly on only need to be included in sufficient numbers in the training set. We demonstrate this advantage with clb1-6Δ mutants that create filamentous buds.

Despite the importance of *S. cerevisiae* as a model organism, to our knowledge, neither does a gold-standard image and segmentation data set exist for yeast nor a CNN trained on such a set. Training data in the form of manual annotations of cell masks is expensive and labor-intensive to generate, especially if it needs to include mutants, which are important for many laboratories. Furthermore, it is not widely known which of a panoply of artificial neural network architectures is suited best, what the disadvantages of each are, and how they can be mitigated.

Previous work demonstrated that a CNN can segment yeast images better than competing methods under very low light conditions.^12^ However, the training set was focused on the specific challenge of very low light levels. YeastSpotter, a CNN for yeast image segmentation based on the Mask-RCNN architecture, was not trained on yeast images but mostly on human cell nuclei.^13^ Thus, it is not surprising that many images of yeast cells cause it to make mistakes (see ‘Comparison to other methods and benchmarking’). The bright-field images of diploid yeast cells published by Zhang et al.^14^ contain in-focus and substantially out-of-focus cells in the same field of view, with only the in-focus cells segmented; it is unclear how a neural network trained on this data set would detect out-of-focus buds well or segment images that are slightly out of focus as in Fig. 1 accurately. The web resource YIT^15^ contains high-quality bright-field and phase contrast images of wild-type yeast cells but only the cell centers are annotated, not the borders.

Beyond yeast, the approach of DeepCell^16^, which was applied to bacterial and mammalian cells and which inspired ours, has the drawback of requiring an additional fluorescent channel for segmentation, which we seek to avoid. Experiments may need all available fluorescent channels for measurements or may need to avoid exciting optogenetic constructs.

Here, we present a large, diverse data set for yeast segmentation and an easy-to-train CNN, which we call YeaZ (pronounced: y-easy). In addition, we have created a Python-based graphical user interface (GUI) to apply the CNN to images in a user-friendly manner, visualize the images and the segmentation masks, apply the bipartite matching algorithm for tracking, and correct potential mistakes. To avoid the need for fluorescent nuclei marking the cell interiors as in the DeepCell method^16^, we seed cells based on peaks of the distance transform and perform a ‘cell-cell boundary test’ to remove erroneous borders. Using the YeaZ CNN to measure the cell geometry of hundreds of wild-type and cyclin mutant cells, we found differences in elongation which indicate that the mitotic cyclin CLB2 controls cell morphology unexpectedly early and gradually. To assess the suitability of the YeaZ CNN without installing any software, images can be submitted to a website for segmentation, accessible under www.quantsysbio.com. Users are invited to submit challenging images for inclusion in the training set, which thus will expand with time and improve the CNN.

## Results

### Data set

We segmented >8400 budding yeast cells of strain background W303, recorded by phase contrast microscopy, semi-manually using a custom image processing pipeline (Fig. 2). In total, this resulted in 372 images and corresponding manual annotation masks, which were checked by 1-2 other people. The set includes normally growing, pre-Start (clnΔ) arrested, filamentous G1/S (clb1-6Δ) arrested, metaphase (cdc20Δ) arrested, and DNA damaged cells, some of which are shown in Fig. 1. Cells were often in large colonies, in which even by eye, cell borders can be difficult to ascertain. Older and bigger cells contained large, transparent inclusions, likely vacuoles, which many classical image segmentation techniques fail to ignore because their edges look like cell borders. Potentially sick cells with strange visible features were included (Fig. 1). Cell sizes varied widely from about 0.4 *μ*m^2^ to 80 *μ*m^2^ (mean wild-type size ≈16 *μ*m^2^). We annotated barely visible buds. Some images were sufficiently out of focus for cells to develop a second light ring around them, which makes identification of the cell edge difficult for many methods (Fig. 1).

**Fig. 2:**
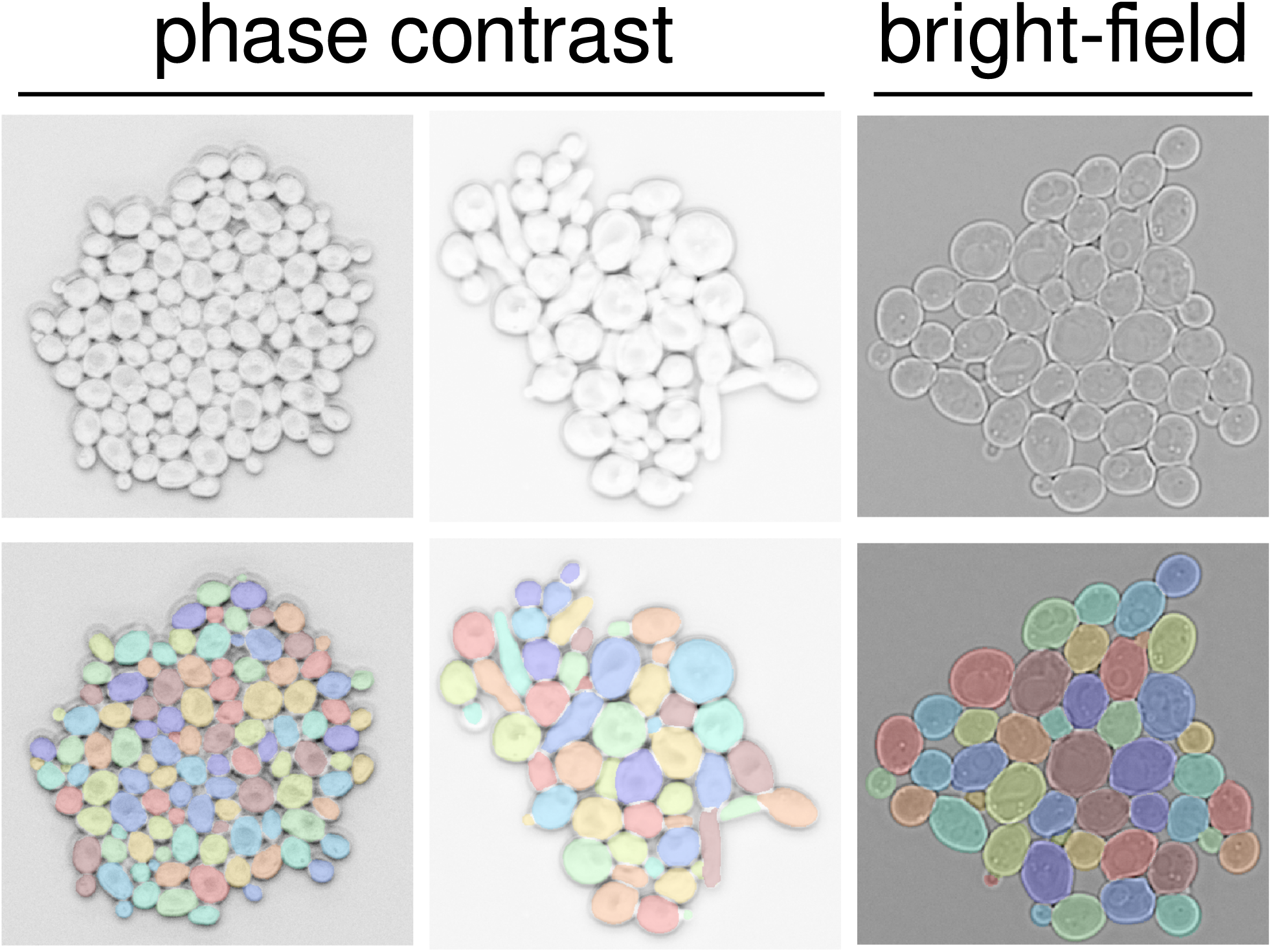
Overview of the YeaZ training data set. Shown are examples of raw images acquired with phase contrast or bright-field microscopy (upper row) and corresponding manual annotations (lower row). Phase contrast image inverted for better visualization.

Using the following trick, we segmented another >1700 cells recorded by bright-field microscopy (Fig. 2): We took images of the same scene of wild-type cycling cells by bright-field and by phase contrast microscopy in rapid succession. Then, we used the YeaZ CNN to segment the phase contrast images efficiently and transferred the segmentation masks to the bright-field images. However, for the rest of our work, we did not use the bright-field segmentations but are making the data available to the community.

Data augmentation artificially increased the size of the training set even further by rotating, flipping, shearing, enlarging or shrinking, as well as dimming or brightening the images for training the CNN.

### Convolutional neural network (CNN)

We evaluated three well-known convolutional neural network architectures: U-Net^17^, Mask-RCNN^10^, and Stardist^18^. We chose U-Net and trained it to distinguish pixels belonging to cell bodies (mapped to 1) from background or cell-cell border pixels (mapped to 0). We did not further distinguish between background and cell-cell border pixels^16^ because the two were often difficult to discriminate unambiguously during the annotation of the training set and because even without this differentiation the resulting CNN was highly accurate. We decided against Mask-RCNN because of artefactual cut-offs of the identified cell regions, presumably due to the method’s rectangular bounding boxes. Stardist alleviates this problem by approximating cell borders with star-convex polygons. Although such a representation is likely appropriate for the typical round shapes of wild-type cells, it is ill-suited for elongated and filamentous cell shapes such as of clb1-6Δ cells.

### Segmentation steps

The CNN assigns to each pixel a score from 0 (border- or background-like) to 1 (cell-like). These continuous scores are turned into a segmented image by the following steps (Fig. 3):

**Fig. 3:**
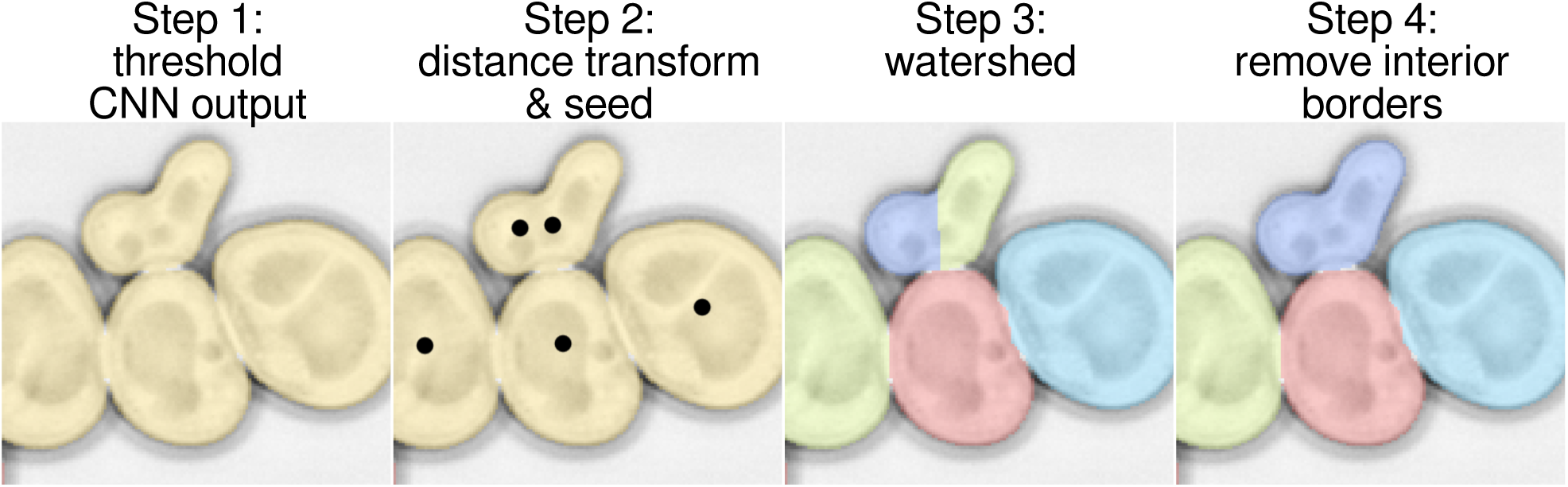
Steps to segmentation: 1) threshold the CNN output, 2) find the peaks of the distance transform (= seeds), 3) watershed, 4) remove erroneous interior borders using a cell-cell boundary test. Phase contrast image inverted for better visualization.

1. *Initial cell-versus-non-cell classification:* A threshold of 0.5, arbitrary but intuitive, is used to distinguish putative cell pixels from the rest. This step already identified most cell bodies as distinct from one another in our images. However, some cells were connected by bridging putative cell pixels, which is why the following steps are needed.
2. *Find a point inside each cell:* For each putative cell pixel, the distance transform (the shortest Euclidean distance to a border/background pixel) is computed. Pixels at which the distance transform has a maximum within a radius of 10 pixels (≈1 *μ*m) identify putative interior points of cells. This step successfully identified one or more points in each cell in our images. (To detect very small buds, we lowered this threshold, see ‘Comparison to other methods and benchmarking’.)
3. *Assign a putative cell to each interior point:* Each peak of the distance transform is used as a seed for the watershed method, which assigns regions of pixels to each peak. These regions are the putative cells.
4. *Remove erroneous cell boundaries:* Since the distance transform may yield more than one point inside each cell, e.g., for a dumb-bell shaped cell, a real cell may be erroneously subdivided into multiple regions by the watershed procedure. This is a well-recognized problem in image segmentation^11^ and could be circumvented, for example, by a fluorescent nuclear marker specifying a unique interior point. To avoid the requirement for an additional channel, we devised the following cell-cell boundary test: For all pairs of putative cells, we evaluate whether the pixels on the boundary are too cell-like; if the average CNN score for the top 3/4 of boundary pixels (bottom 1/4 is ignored because an erroneous boundaries will touch real boundaries at their two ends) is above 0.99, i.e., very cell-like, this boundary is likely erroneously subdividing a real cell. In that case, the two regions separated by the erroneous boundary are merged. This strategy fixed all cases of split cells that we encountered, which, for example, occurred for 10% of cells in Fig. 4 (top). We did not observe that any cells were joined erroneously.

**Fig. 4:**
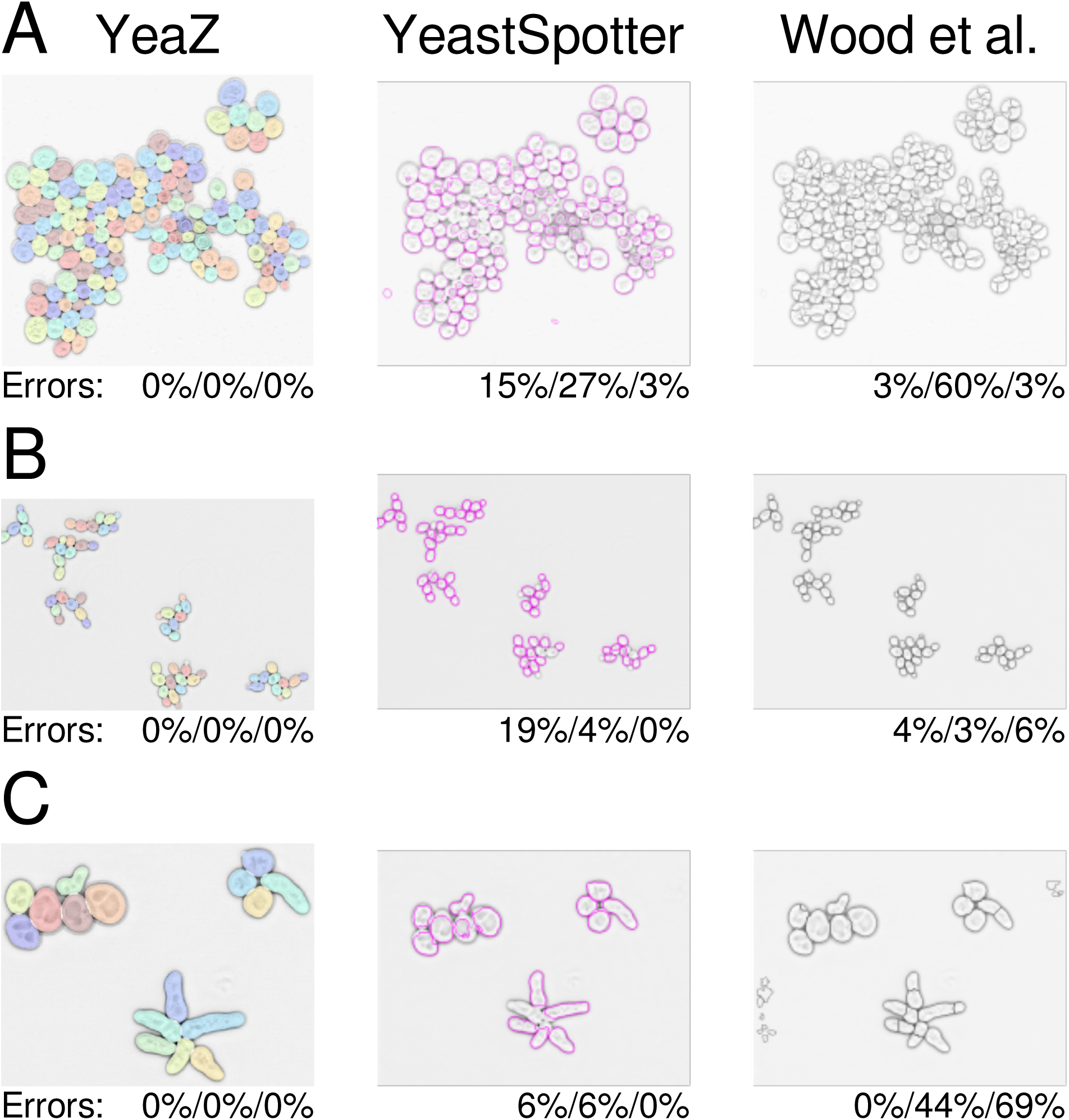
Comparison of YeaZ with YeastSpotter and Wood et al. ^8^ The error values represent the fraction of missed cells / of bad contours / and of spurious cells. Phase contrast images are inverted for better visualization.

We introduced a small number of parameters in the above steps without fine-tuning because the results did not require it (see ‘Comparison to other methods and benchmarking’).

### Tracking

The tracking algorithm is similar to the one in CellStar^7^. Cells are matched between two consecutive frames. For each time point *t* and each cell *i*, the center of mass and the area are calculated (*x*_*i*_ (*t*), *y*_*i*_ (*t*), *A*_*i*_ (*t*). The mean over all cells at time *t* is subtracted and the resulting 3-tuples are rescaled to normalize the variances to (3, 3, 1. (We observed that weighting in favor of the position makes the algorithm work better.) The actual tracking step is performed by bipartite graph matching, finding pairings that minimize the summed Euclidean distances between the normalized 3-tuples. This can be done efficiently by the Hungarian algorithm^19,20^. We have found the method to work with very few errors.

### Comparison to other methods and benchmarking

The convolutional neural network approach has at least two inherent advantages over non-machine learning approaches: While diverse and potentially difficult to analyze, budding yeast cells may only have a range of shapes and visible features. Our large, diverse data set is well-suited to cover this range and enables the neural network to interpolate between shapes it has already been trained with in order to segment new images. Furthermore, should a particular condition or cell type not yield satisfactory segmentation results, the addition of a number of new examples in principle suffices to expand the capabilities of the neural network, as we demonstrate for clb1-6Δ cells.

Ideally, to compare YeaZ to other methods, we would use a gold-standard segmentation benchmark. However, we could not find such a data set, which is in part why we believe our data set of segmented images will be useful to the community. Instead, we proceed as in a comprehensive comparison of segmentation methods performed previously ^15^. We begin by focusing on three images: one image of moderate complexity from our timelapse recordings that was not included in the training set (A) and two images of budding yeast cells included in the prior comparison^15^ containing cycling (B) or pheromone-arrested (C) wild-type cells, respectively. Images B and C represent the last time points in two timelapse recordings (data sets 9 and 10 in ref.^15^) and thus are the most complex images of the series, containing the largest number of cells. Together, the three images cover three important situations: a crowded scene (A), a relatively sparse scene (B), and new shapes (C) not included in our training set.

We chose for our comparison the newest segmentation method we found published, by Wood et al.^8^, and the only other available neural network for yeast segmentation YeastSpotter^13^, which was, however, not trained on yeast cell images. Wood et al.’s method has at least 16 parameters; we varied min_cell_size, max_cell_size, min_colony, clean_BW, and size_strel_bg_2 away from from the default values to improve the results. Since the method by Wood et al.^8^ compares favorably with other published methods^15^ and given the stark differences in the segmentation qualities we observed, we believe that confining ourselves to two reasonable comparisons suffices.

The results of the YeaZ segmentation are perfectly accurate for all three images (Fig. 4). No tweaking of our parameters was necessary except to adjust roughly the pixel equivalent of the 1 *μ*m threshold for the distance transform to the larger pixel sizes in B and C. Close inspection of the results revealed no missed cells, no missed buds, no errors in the boundary assignments, and no false cells. Given the many differences between our strains and conditions and those of the images from ref.^15^, this exemplifies the transferability of the YeaZ CNN.

In order to compare the results in a way that is useful for the typical user, we scored the output of the other methods by counting the number of cells that were missed by the segmentation, that were segmented clearly badly, or that were likely acceptable for most purposes (Fig. 4). The other methods’ boundaries were not required to be perfect; we scored their output rather leniently. Our detailed scoring is presented in Figs. S1-S6.

The scene with widely varying cell sizes caused both YeastSpotter and Wood et al.’s method to make many mistakes (Fig. 4 top row). Generally, Wood et al.’s method tended to overseg- ment, i.e., subdivide cells erroneously. YeastSpotter tended to miss cells.

To find out how low the error rate of the YeaZ CNN may be, we analyzed the entire data sets 9 and 10 from ref.^15^. The resulting segmentations were flawless for data set 9 for all 1596 cells except for four buds; tiny buds of a few pixels were detected early except for four (Fig. S7), which were detected at the next time point when they were slightly bigger. The error rate is thus 0.25%. For data set 10, all 484 cells were segmented accurately (error rate: 0%), however, we remark that the images in data set 10 are very similar to each other.

Thus, on images from us and others that are challenging for other methods, YeaZ produced ground-truth level segmentations.

To complement this analysis with a mathematically rigorous comparison, we also scored all three methods, YeaZ, YeastSpotter, and Wood et al., computationally. We took 17 semi-manually segmented phase contrast images containing 1894 wild-type cycling cells, which were not included in the training set for the YeaZ CNN, and computed standard segmentation metrics such as accuracy and mean intersection-over-union (IoU)^18^ (Fig. 5). The YeaZ CNN performed very well (mean accuracy: 94%) with most of the missed cells being small buds that the CNN delimited differently than the human annotators. Given that many of these buds spanned only a few pixels (see Fig. S7 for examples of small buds), it was easy for two slightly different segmentations to differ by the 50%, threshold for the accuracy metric – without the bud actually having been missed or clearly incorrectly segmented. By both metrics, YeastSpotter showed a substantially higher error rate than the other methods. Wood et al.’s method performed better than YeastSpotter on this set of images (mean accuracy: 79%). (Similarly, among the three test images in Fig. 4, Wood et al. had performed reasonably well for wild-type cycling cells (middle row).)

**Fig. 5:**
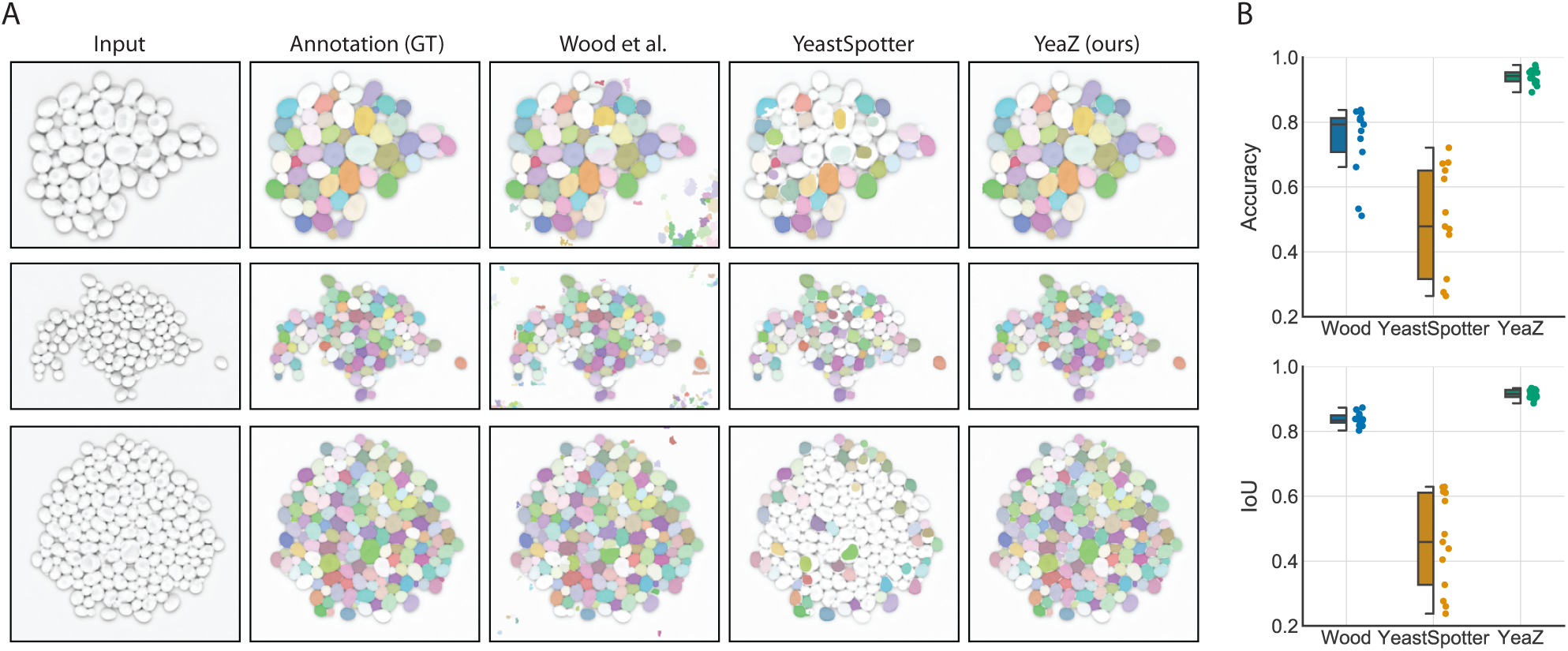
Detailed computational comparison of all methods on test images of cycling wild-type cells not included in the YeaZ training set. A) Each row shows an example test image, its ground-truth annotation (GT), and the result of Wood et al. ^8^, YeastSpotter ^13^, and YeaZ, respectively. B) Quantification of segmentation performance of all methods. As is common in the computer vision literature, we call a predicted cell a true positive (TP), if its intersection over union (IoU) with the corresponding ground-truth (GT) cell is larger or equal than 50%. Similarly false positives (FP) and false negatives (FN) are defined as predictions that have no GT match and vice versa. As segmentation metric we show average accuracy 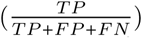 and average intersection-over-union of true positives (*IoU*). Boxes show interquartile range (IQR), lines signify median, and whiskers extend to 1.5 IQR.

### Expanding the capabilities of the CNN

In order to gauge the adaptability of the CNN to new cell shapes, we trained it with and without approximately 50 filamentous clb1-6Δ cells growing in different colonies. We then tested the CNN on another image from a later timepoint of one of the scenes, when the filamentous cells had grown substantially longer (Fig. 6). Importantly, these longer cells were not broken up by the CNN trained on the expanded data set. Note that these colonies can be very difficult to segment by eye in the places where cells are crowded; thus, the mistakes that are made when strangely shaped mutant cells surround and partially overlap with each other as in the bulk in Fig. 6 may be expected, given the number of clb1-6Δ mutants in the training set.

**Fig. 6:**
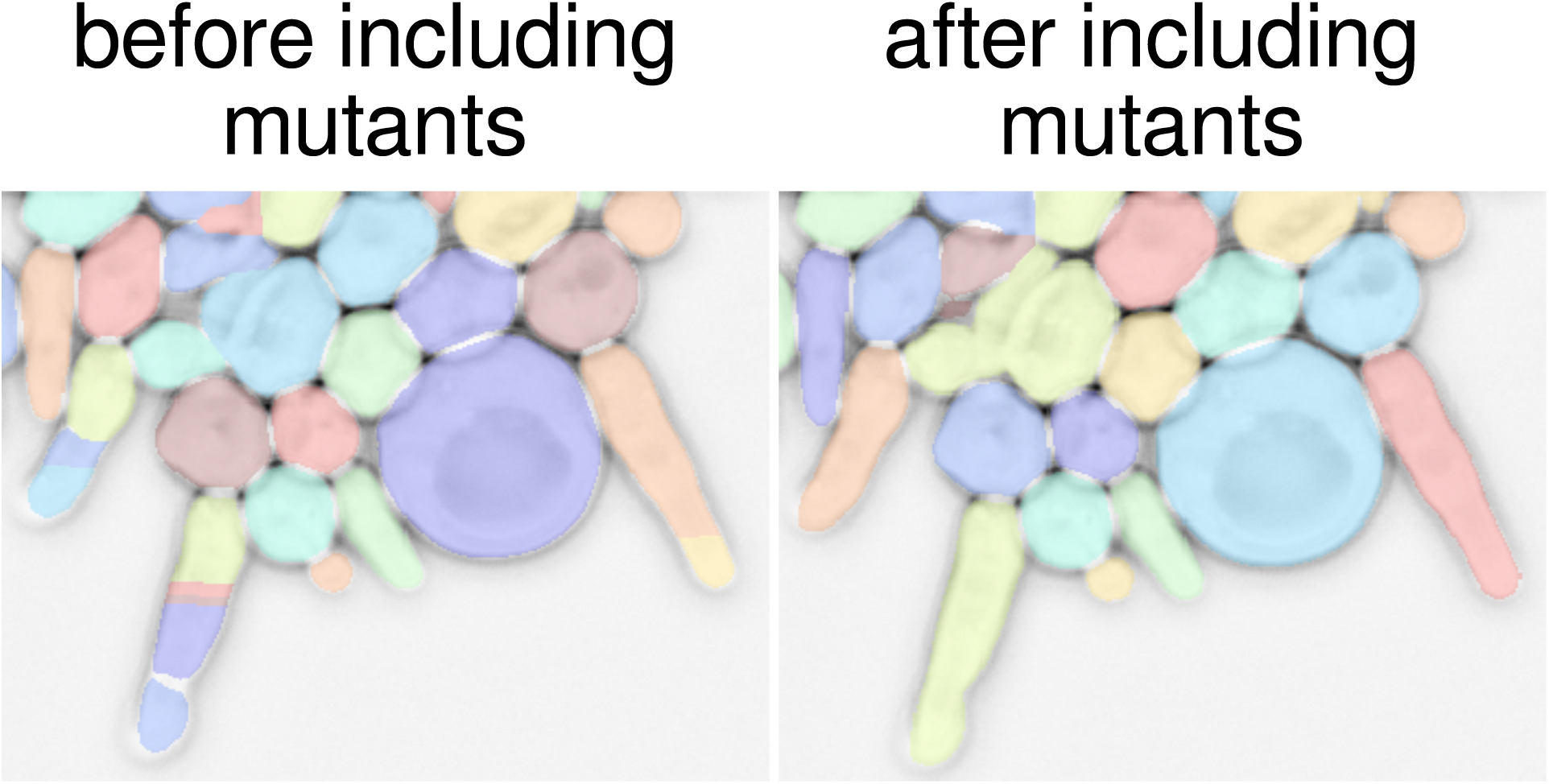
Adaptability of the CNN. clb1-6Δ mutants were either excluded (left) or included (right) in the training set for the CNN, which was then tested on an image of clb1-6Δ cells from a later time point with even longer filaments, shown here. Note that the color of each cell is dependent on the internal numbering and therefore arbitrary. However, there are no fragmented filamentous cells on the right although there are segmentation errors when the strangely shaped cells are crowded. Phase contrast image inverted for better visualization.

One solution to minimize the manual labor required to expand the training set is to proceed iteratively: segment a few images under a new condition, retrain the neural network with these images, and repeat with an improved neural network until the performance is acceptable.

### Graphical user interface (GUI)

To apply the CNN and the tracking algorithm and correct their mistakes, we designed a Python-based graphical user interface (GUI) (Fig. 7). New cells can be drawn, modified after segmenting with the CNN, and cells can be relabeled. We were inspired by Microsoft Paint to include image manipulation tools such as brushes and erasers to manipulate the segmentation masks. The user can leaf through timelapse images with the current, the previous, and the next time point shown simultaneously, which can be helpful for verifying small buds. Fields of view and imaging channels can be changed. The GUI can read in multi-layer TIF files, folders of multiple TIF files, and Nikon ND2 files. We are continuously improving the capabilities of the GUI since we are using it ourselves. The latest version can be found through our website http://www.quantsysbio.com.

**Fig. 7:**
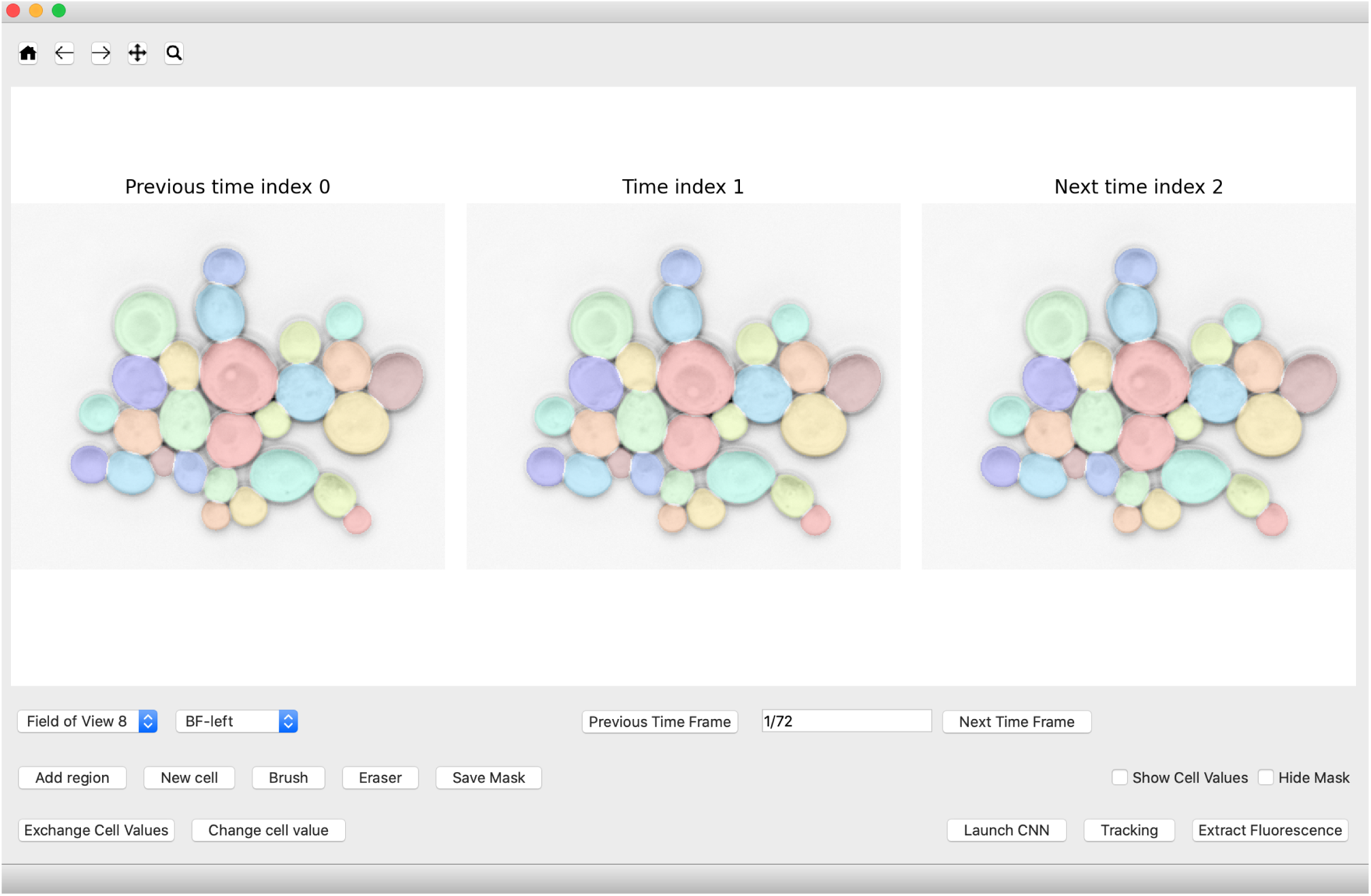
GUI for applying the YeaZ CNN, tracking cells, and correcting segmentations.

### Cell shape measurements reveal the timing and strength of morphogenesis control

New yeast daughter cells grow as buds from the tip until mitotic cyclins, mainly Clb2, change the direction of growth from apical to isotropic (Fig. 8 A); overexpressing the cell cycle Start initiator CLN2 or deleting CLB2 leads to more elongated cells.^21,22^ Since Clb2 turns on as part of a positive feedback loop some time after cell cycle Start,^23^ one may expect growth depolarization to occur suddenly at a specific time after budding. To investigate when this switch occurs, we analyzed the geometries of hundreds of wild-type and mutant cells using YeaZ (Fig. 8 B, C). We quantified each cell’s elongation (=major axis/minor axis) by equating its second moments of the area with those of an ellipse. We used the cells’ areas as stand-ins for the time after budding because cells grow in size continuously.

**Fig. 8:**
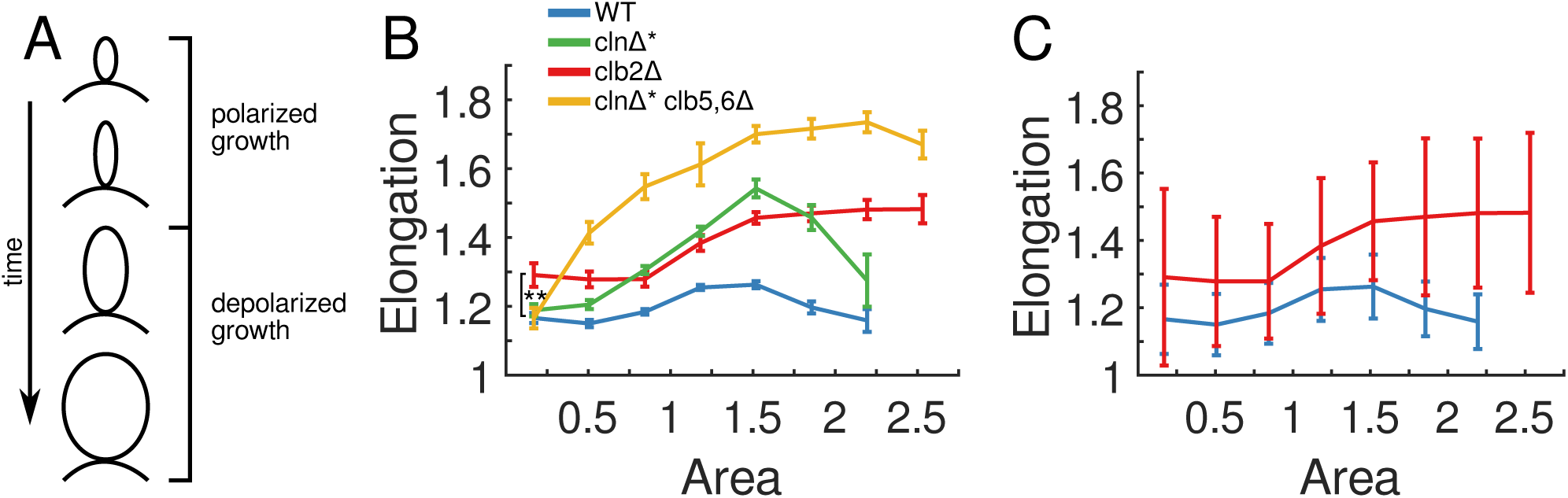
Clb2 promotes a more circular cell shape beginning early in the cell cycle. A: Schematic of cell shape during growth. B: Mean and SEM of elongation versus area for different cyclin mutants. **: p <0.01, single-tail est. C: Mean and STD for two of the populations from panel B illustrating the variability in the data. B, C: Abscissa scaled so that 1 is the mean area of wild-type cells, corresponding to 16.3 *μ*m^2^. n=525 (WT), 500 (clnΔ*), 597 (clb2Δ), 543 (clnΔ* clb5,6Δ). The largest 2.5% of the overall population was discarded. The remaining range was binned into 8 equal intervals. There were no WT or clnΔ* cells in the largest bin, therefore, no values are shown.

Wild-type cells (blue) initially became more elongated with size, and depolarization kicked in when cells reached around 1-1.5 of the mean wild-type area, making cells more circular again (Fig. 8 B).

cln1-3Δ MET3pr-CLN2 (abbreviated: clnΔ*) cells (green) which expressed the Start cyclin CLN2 continuously in medium lacking methionine started as buds that were similarly shaped as wild type but became substantially more elongated (Fig. 8 B). Subsequently, however, they depolarized and grew sufficiently to end up about as circular as wild-type cells when they were large.

Interestingly, clb2Δ cells (red), were already substantially more elongated when they were very small (first bin: 0-33% of mean wild-type area), even though Clb2 seemed to depolarize wild-type cells much later at around 1-1.5 mean wild-type area (compare wild-type and clb2Δ in Fig. 8 B). Thus, Clb2 influenced cell morphology already very early in the cell cycle, potentially because of i) early, weak activation of CLB2, ii) basal expression of CLB2, or iii) left-over Clb2 from the previous cycle. Furthermore, clb2Δ cells could not be detected to depolarize at all, and became even more rod-like with size.

To test whether Clb2 is initiated at low levels earlier than previously thought, we analyzed clnΔ^∗^ clb5,6Δ (yellow) cells (Fig. 8 B). CLB5 and CLB6 are key activators of CLB2. ^23^ We combined their deletions with the clnΔ^∗^ mutations and constructs, which produce Cln2 continuously in minus methionine medium. This was done to maximize polarized growth, which Cln2 promotes, and compensate for the loss of Clb5, which might also promote polarized growth as it can substitute for cell cycle Start initiators. Nevertheless, cells started similarly shaped as wild-type or clnΔ^∗^ cells, inconsistent with explanation i). Explanations ii) and iii) would be surprising because Clb2 is known to interfere with origin of replication licensing in G1 phase^24^, before cell cycle Start; however, ii) or iii) would be consistent with the requirements for origin of replication licensing if licensing is less sensitive to Clb2 than morphogenesis control.

In summary, our data refine our understanding of the timing and strength of morphogenesis control, to be earlier than commonly thought and to be gradually strengthening with time, not simply switching on-off. Both results are surprising considering the known timing and manner of activation of Clb2. Since we focused primarily on the geometry of the smallest cells, any potential minor differences in growth rates between the mutants should not affect our conclusions.

This application exemplifies why an efficient segmentation method is needed and how it can provide new insights. Because variability is high (see standard deviation in Fig. 8 C), large numbers of cells are needed for statistically significant results (see standard error of the mean in Fig. 8 B).

## Discussion

We present the first freely available, large, diverse, and high-quality set of segmented yeast images as well as a CNN trained on this data set. The CNN segments new images recorded by us and others very accurately. We introduced a simple cell-cell boundary test to alleviate the oversegmentation problem that arises in the absence of an established unique interior point, which a fluorescent nuclear marker provides in other methods^16^. Our approach does not require extra fluorescent markers.

We applied the CNN to detect that, likely, basal CLB2 expression or left-over Clb2 from the previous cell cycle influence cell shape early in a new cycle, not just when cells depolarize markedly. This is surprising and, to our knowledge, has not been observed previously.

While we designed our data set to be likely sufficiently diverse for most applications, there may arise conditions under which it is not. Should the CNN perform poorly for certain new cell shapes or conditions, in principle, adding challenging semi-automatically segmented training examples to the current set ought to improve the performance, as we demonstrated for clb1-6Δ cells, or perfect it. Repeated cycles of segmenting with an incrementally improving CNN, correcting mistakes, and retraining may be a particularly labor-efficient way to expand the capabilities of the CNN.

As a proof of principle for how the existing CNN can be leveraged to improve it further, we applied a simple trick to expand the training set beyond phase contrast images: We recorded the same scene with both phase contrast and bright-field microscopy and used the CNN to segment the phase contrast images. This gave us a training set for bright-field images with little effort. We make the bright-field segmentations available although we did not train the CNN with it.

## Methods

### Microscopy

Images were recorded of cells growing in microfluidic chips with a 60x objective and a Hamamatsu Orca-Flash4.0 camera.

### Pre-processing

The training set consists of i) microscopy images and ii) mask images from the semi-manual annotation (see ‘Data set’) which are of the same size as the microscopy images and whose pixels denote the ID numbers of the cells in the corresponding microscopy images. Background pixels correspond to 0 in the masks. Before setting all cell numbers to 1 for training the neural network, we found the borders between different cells by dilating each cell and identifying intersecting pixels. These border pixels were then also set to 0 in the mask images. The training set was cut into 256 x 256 images, which overlapped by at least half in width or height, for the training. (The GUI applies the CNN to whole images without cropping.)

### Training

We downloaded the U-net implementation from https://github.com/zhixuhao/unet and adapted it. Batch sizes were set to 25 and training was carried out for 100 epochs. Augmentation was performed with rotation range 90 degrees, shear range 45 degrees, zoom range 0.5-2, horizontal and vertical flipping, and brightness range 0.5-1.5.

### Strains

All strains were W303 based. All except AS18 have been characterized previously.^25,26^

| Name | Mating type<br>locus | Genotype |
| --- | --- | --- |
| AX42-25A | MATa | wild type W303 |
| JB204-6B | MATa | ADE2 HTB2-mCherry::HIS5 |
| SJR112a12d | MATα | cln1Δ0 cln2Δ0::MET3pr-CLN2 cln3Δ0 HTB2-mCherry::HIS5 ura3Δ0 |
| SJR33h6d | | clb2Δ::LEU2 trp1Δ::TRP1::GAL-SIC1(1X) HTB2-mCherry:HIS5 MYO1-GFP::KanMX |
| SJR36a11b | MATa | cln1Δ cln2Δ:CLN2pr-Venus:TRP1 cln3Δ:LEU2 trp1Δ:TRP1::MET3pr-CLN2 clb5Δ:HIS3 clb6Δ:KanMX HTB2-mCherry:HIS5 |
| CL86-1-11D | MATa | clb1Δ clb2Δ::GALL-CLB2::URA3 clb3Δ::TRP1 clb4Δ::his3Δ::kanMX clb5Δ::kanMX clb6Δ::kanMX swe1Δ::TRP1 ADE2 |
| 2147-1C | MATα | cdc20Δ::LEU2 ADE2::GALL-CDC20 |
| AS18 | MATαsyn | cln1Δ0 cln2Δ0::MET3pr-CLN2 cln3Δ0::GAL1pr-HO HTB2-mCherry::HIS5 ADH1pr-HOcs-YFP |

## Software and data availability

The latest version of the YeaZ GUI and data sets can be downloaded through our website, where the CNN can also be tested online:

http://www.quantsysbio.com

## Author contributions

AME, DAHLJ, VG, AS, CHMC, and SJR segmented images. ND, MM, and SJR trained the neural network. ND, MM, and SJR designed the GUI. DAHLJ designed the web interface. VG performed experiments and analyzed the measurements. MM, VG, MW, AFB, and SJR wrote the manuscript. MK gave technical advice. SJR supervised the work.

## Supplementary Figures

**Fig. S1:**
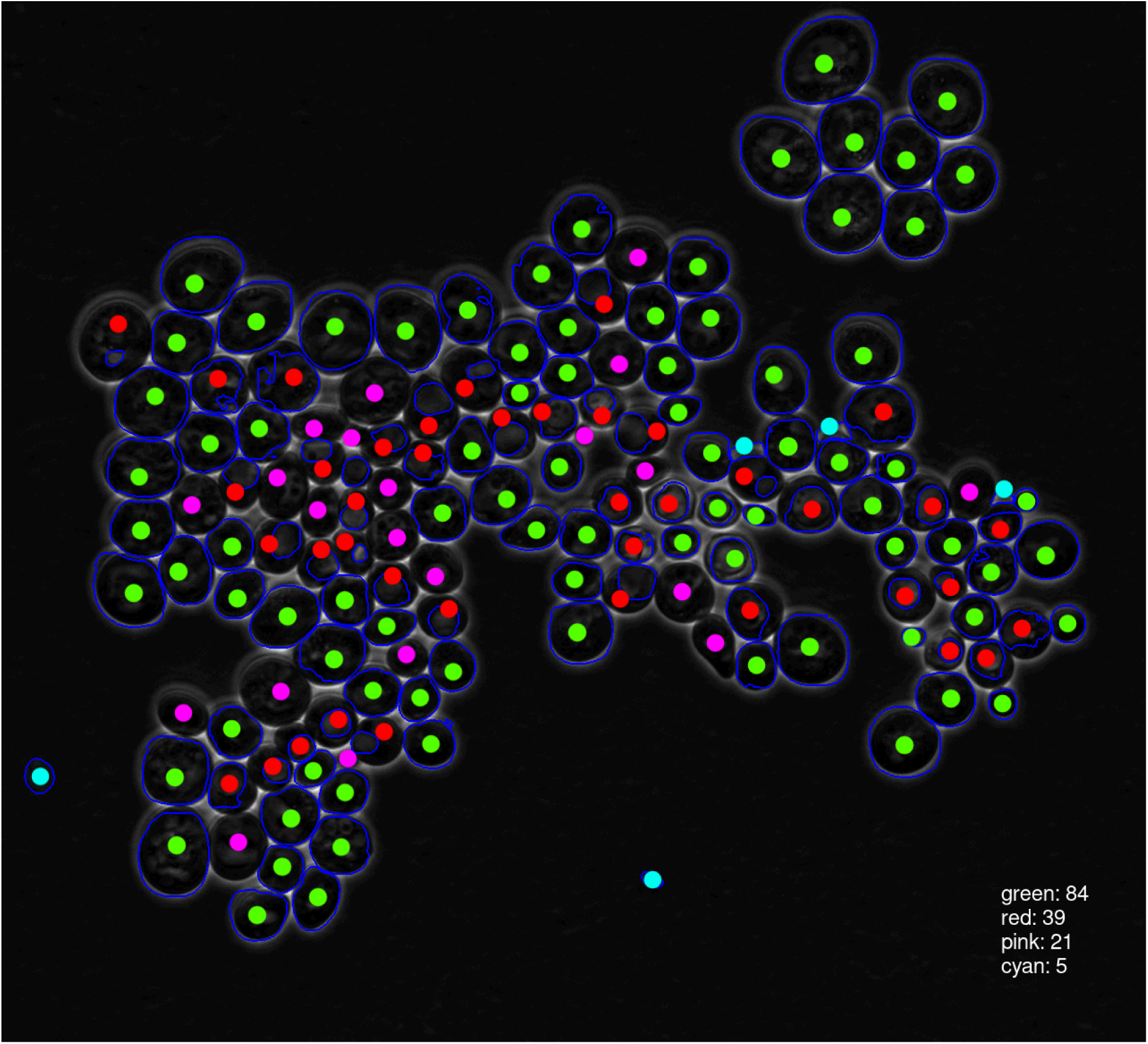
Scoring of YeastSpotter results. Green dots: acceptable segmentation, red: bad segmentation, pink: missed cell, cyan: spurious cell.

**Fig. S2:**
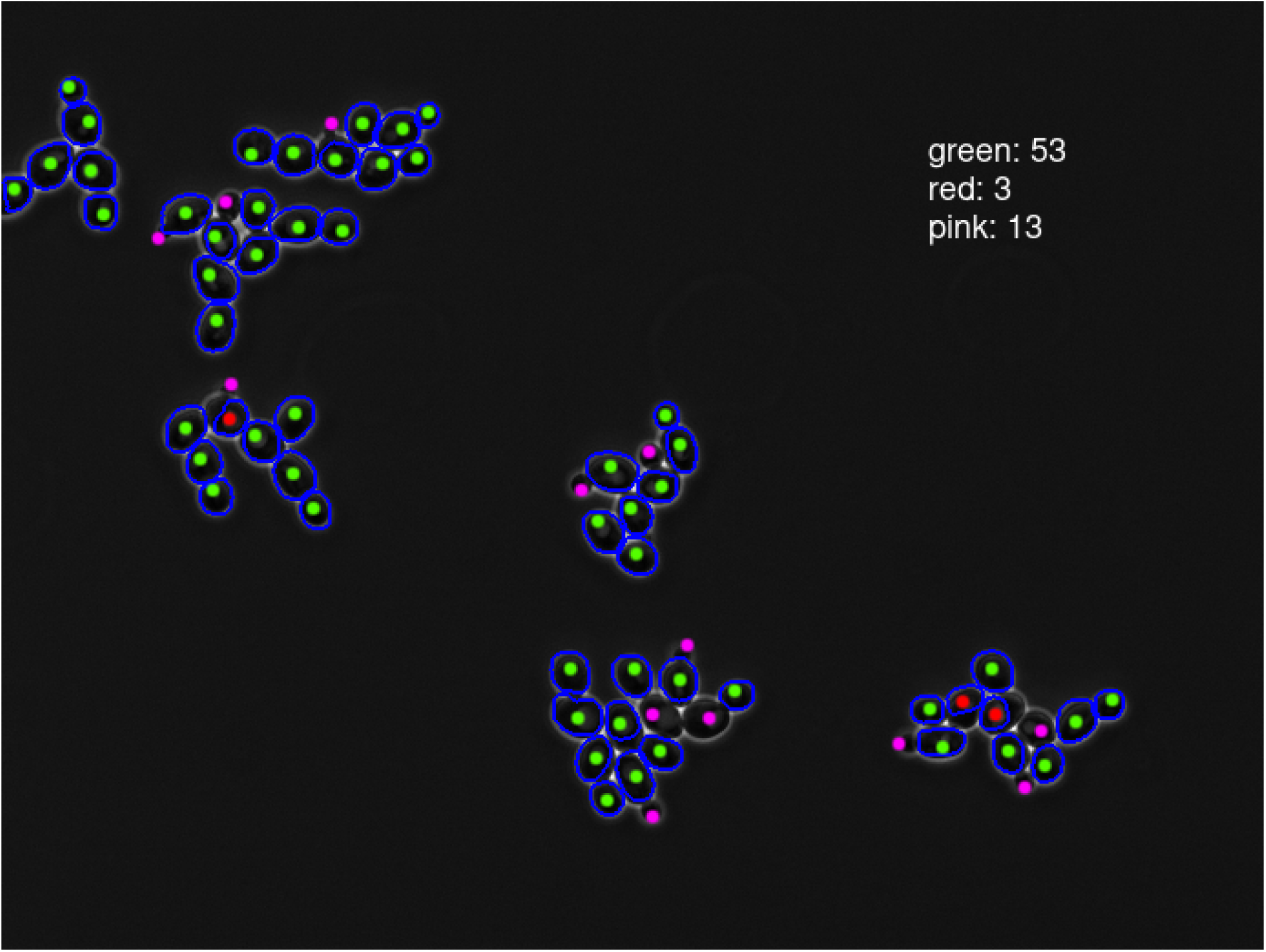
Scoring of YeastSpotter results. Green dots: acceptable segmentation, red: bad segmentation, pink: missed cell, cyan: spurious cell.

**Fig. S3:**
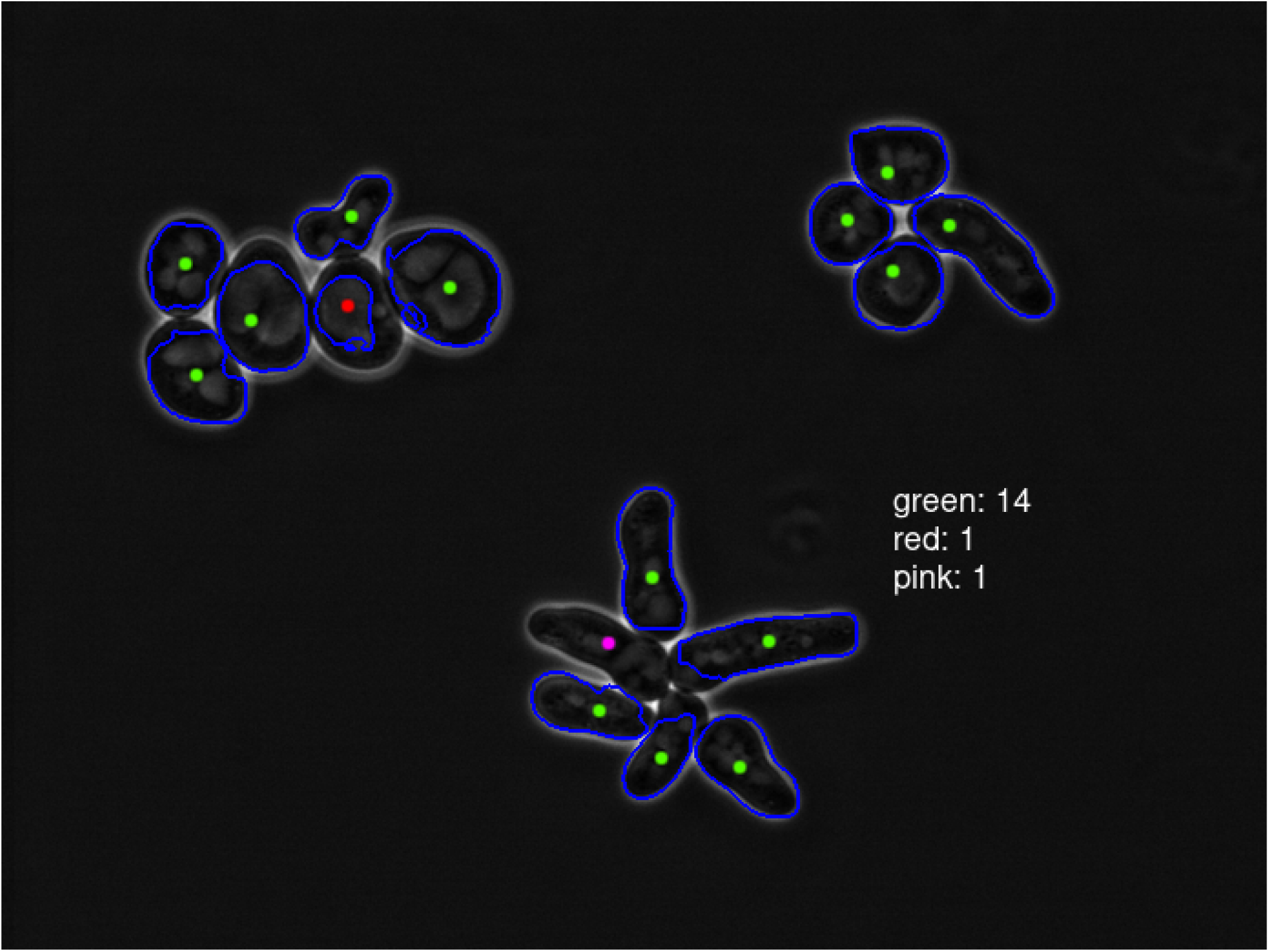
Scoring of YeastSpotter results. Green dots: acceptable segmentation, red: bad segmentation, pink: missed cell, cyan: spurious cell.

**Fig. S4:**
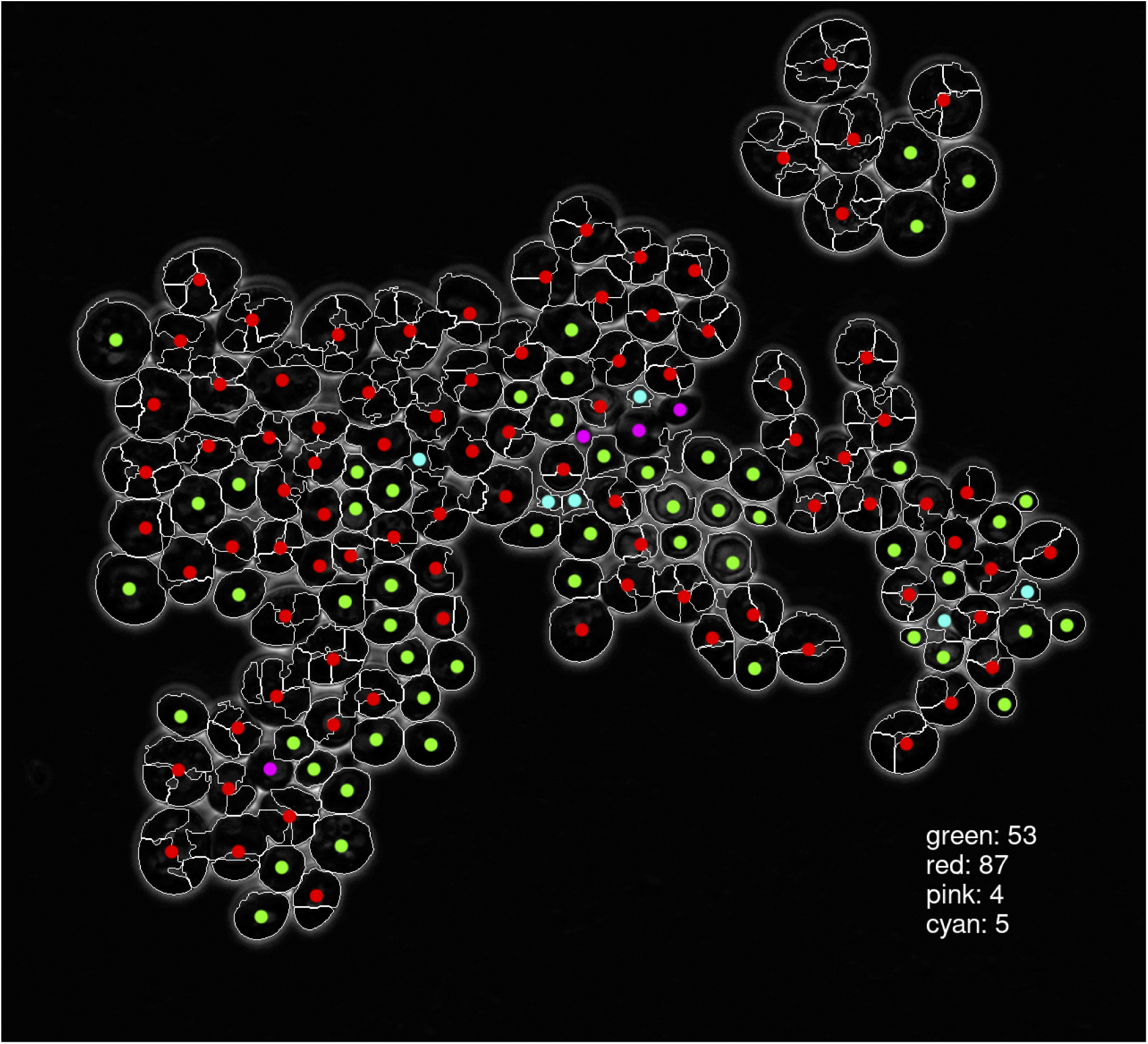
Scoring of the results of the method by Wood et al. ^8^. Green dots: acceptable segmentation, red: bad segmentation, pink: missed cell, cyan: spurious cell.

**Fig. S5:**
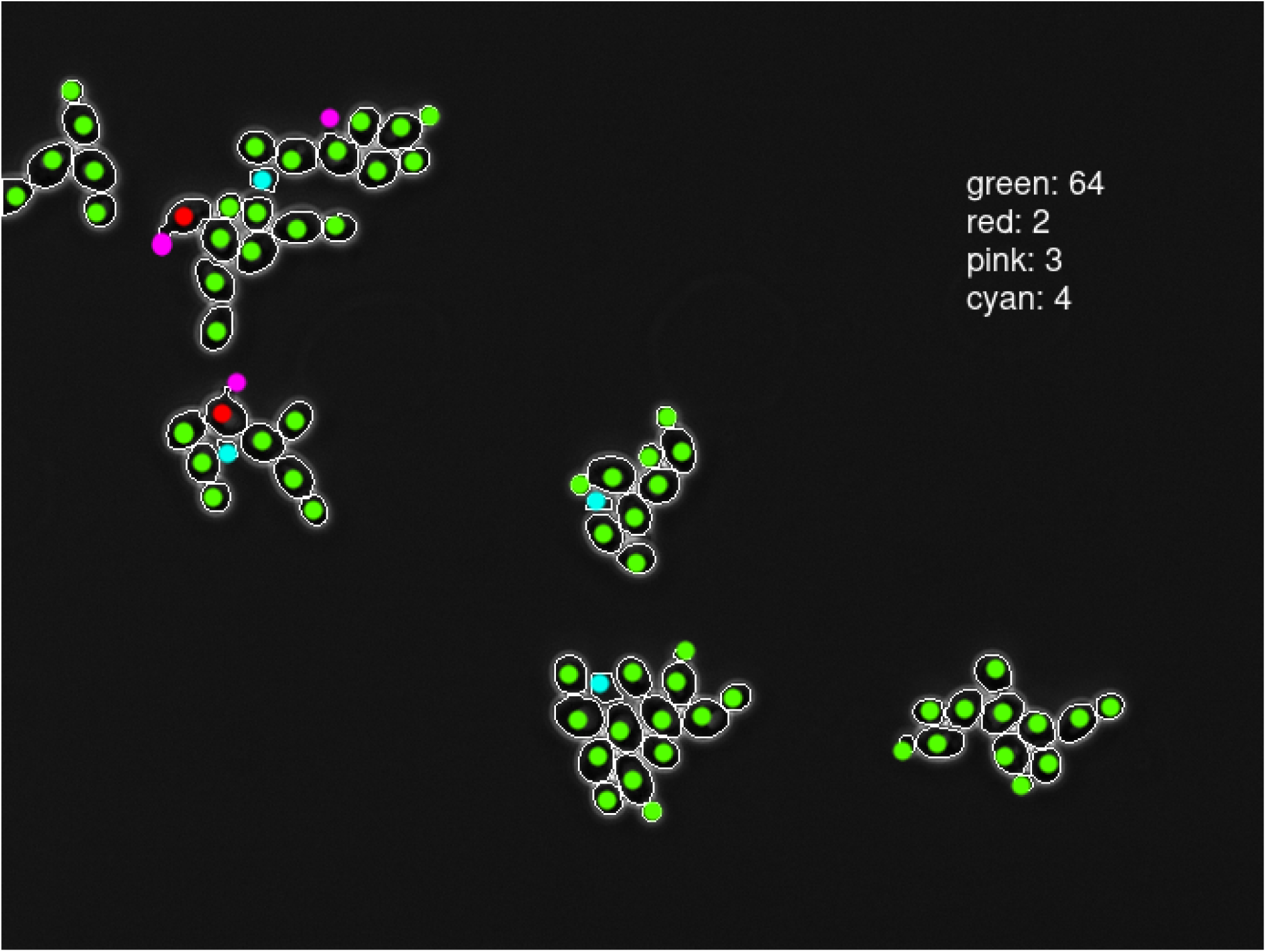
Scoring of the results of the method by Wood et al. ^8^. Green dots: acceptable segmentation, red: bad segmentation, pink: missed cell, cyan: spurious cell.

**Fig. S6:**
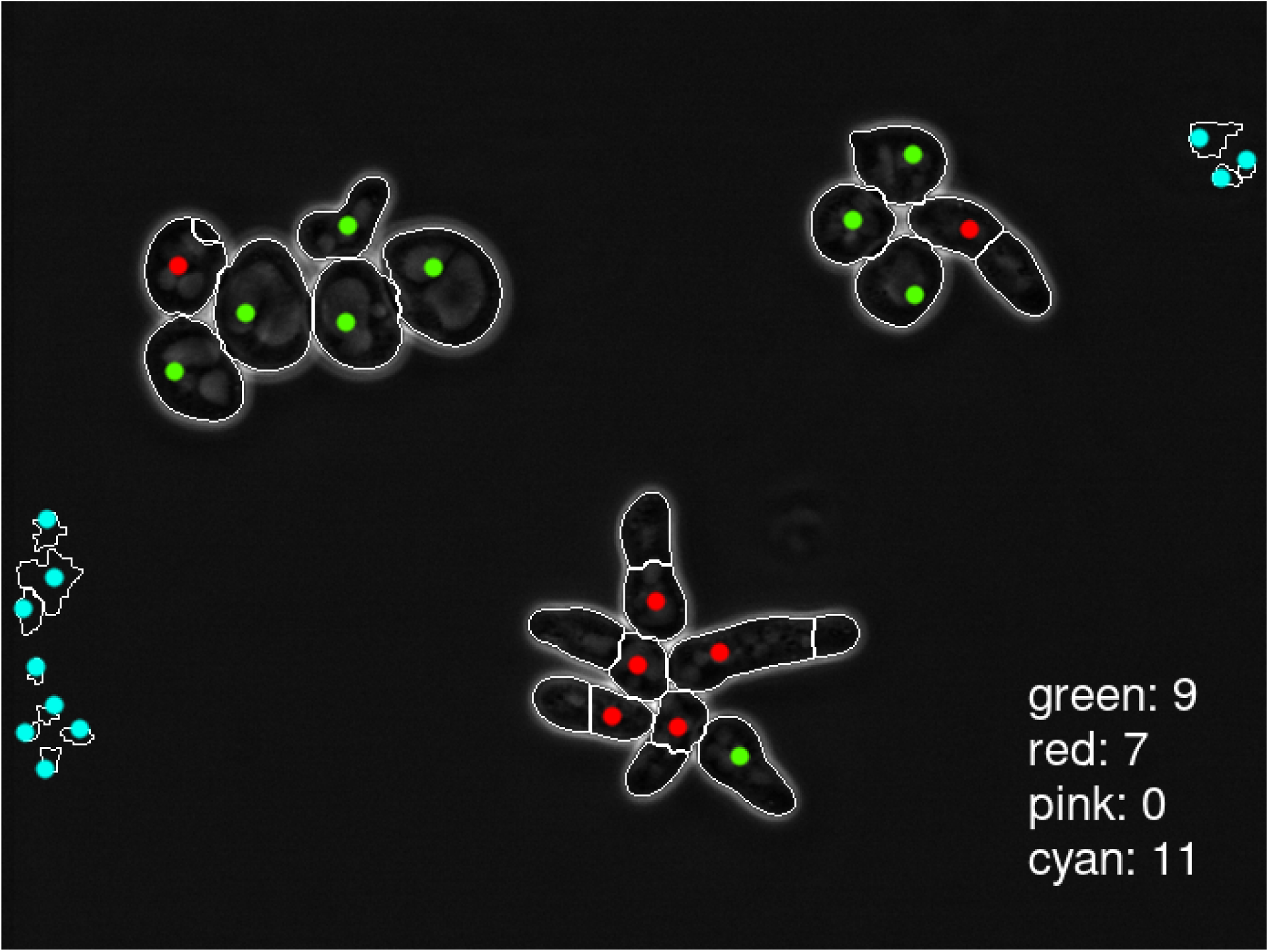
Scoring of the results of the method by Wood et al. ^8^. Green dots: acceptable segmentation, red: bad segmentation, pink: missed cell, cyan: spurious cell.

**Fig. S7:**
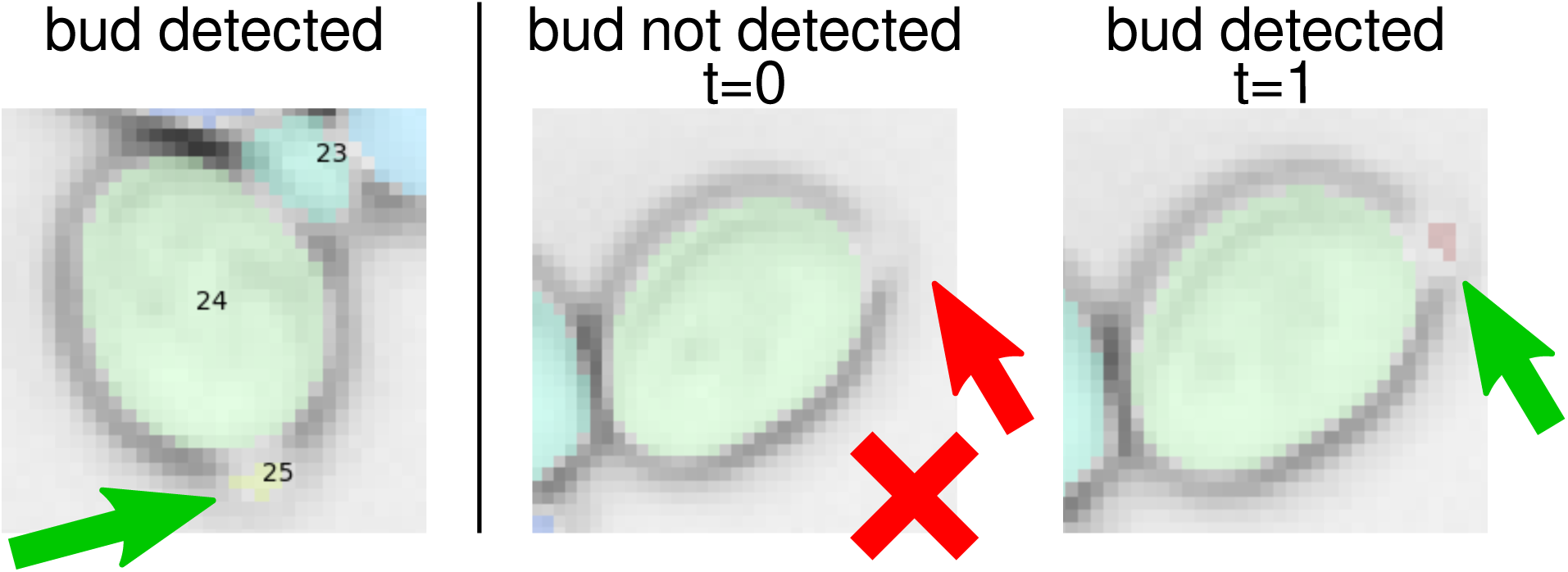
Examples of tiny buds in data set 9 from ref. ^15^, some only a few pixels in size, that were detected by the YeaZ CNN (left and right) except for four such tiny buds (middle example). The missed bud (middle) was, however, detected at the next time point in the timelapse series (right) when it was slightly bigger.

